# CETP expression in females increases body metabolism under both cold exposure and thermoneutrality contributing to a leaner phenotype

**DOI:** 10.1101/2024.11.11.623058

**Authors:** Júlia Z. Castelli, Helena F. Raposo, Claudia D. C. Navarro, Carolina M. Lazaro, Marina R. Sartori, Ana Paula Dalla Costa, Pedro A. S. Nogueira, Lício A. Velloso, Anibal E. Vercesi, Helena C. F. Oliveira

**Affiliations:** Dept of Structural and Functional Biology, Biology Institute, State University of Campinas, Campinas, SP, Brazil; Obesity and Comorbidity Research Center (OCRC), State University of Campinas, Campinas, SP, Brazil; Dept of Pathology, Faculty of Medical Sciences, State University of Campinas, Campinas, SP, Brazil

**Keywords:** CETP, cold exposure, thermoneutrality, body composition, brown adipose tissue, respiration, RRID:IMSR_JAX:003904, RRID:IMSR_JAX:001929, RRID:MGI:7264953

## Abstract

Susceptibility to obesity differs depending on the genetic background and housing temperatures. We have recently reported that CETP expressing female mice are leaner due to increased lipolysis, brown adipose tissue (BAT) activity and body energy expenditure compared to non-transgenic (NTg) littermates under standard housing temperature (22°C). The aim of this study is to evaluate how CETP expression affect body temperature, composition and metabolism during cold exposure (4°C) and thermoneutrality (30°C). When submitted to cold, CETP mice maintained rectal temperature, body weight and food intake similarly to NTg mice along acute or chronic exposure to 4ºC. The body oxygen consumption in response to an isoproterenol challenge was 21% higher at 22ºC, and 41% higher after 7 days of cold exposure in CETP than in NTg mice. In addition, BAT biopsies from CETP mice showed reduced lipid content and increased basal oxygen consumption rates. Under thermoneutrality (30ºC), when BAT activity is inhibited, CETP mice showed higher rectal and tail temperatures, increased food intake and increased energy expenditure. Lean mass was elevated and fat mass reduced in CETP mice kept at 30ºC. In this thermoneutrality condition, soleus muscle, but not gastrocnemius or liver of CETP mice showed increased mitochondrial respiration rates. These data indicate that CETP expression confers a greater capacity of elevating body metabolic rates at both cold exposure, through BAT activity, and at thermoneutrality, through increased muscle metabolism. Thus, the CETP expression levels in females should be considered as a new influence in the contexts of obesity and metabolic disorders propensity.

**NEW & NOTEWORTHY:** We demonstrate here that CETP expression in females increases body metabolism under cold (4ºC) and thermoneutrality (30ºC). Since this has also been shown at 22ºC, it seems a constitutive feature of CETP expression. Brown adipose tissue and red fiber muscle contribute to the overall high metabolism and leaner phenotype of CETP mice. Elevated mitochondrial respiration rates were demonstrated in these tissues. Thus, CETP is a new relevant variable in the context of obesity and metabolic disorders.

**GRAPHICAL ABSTRACT:** 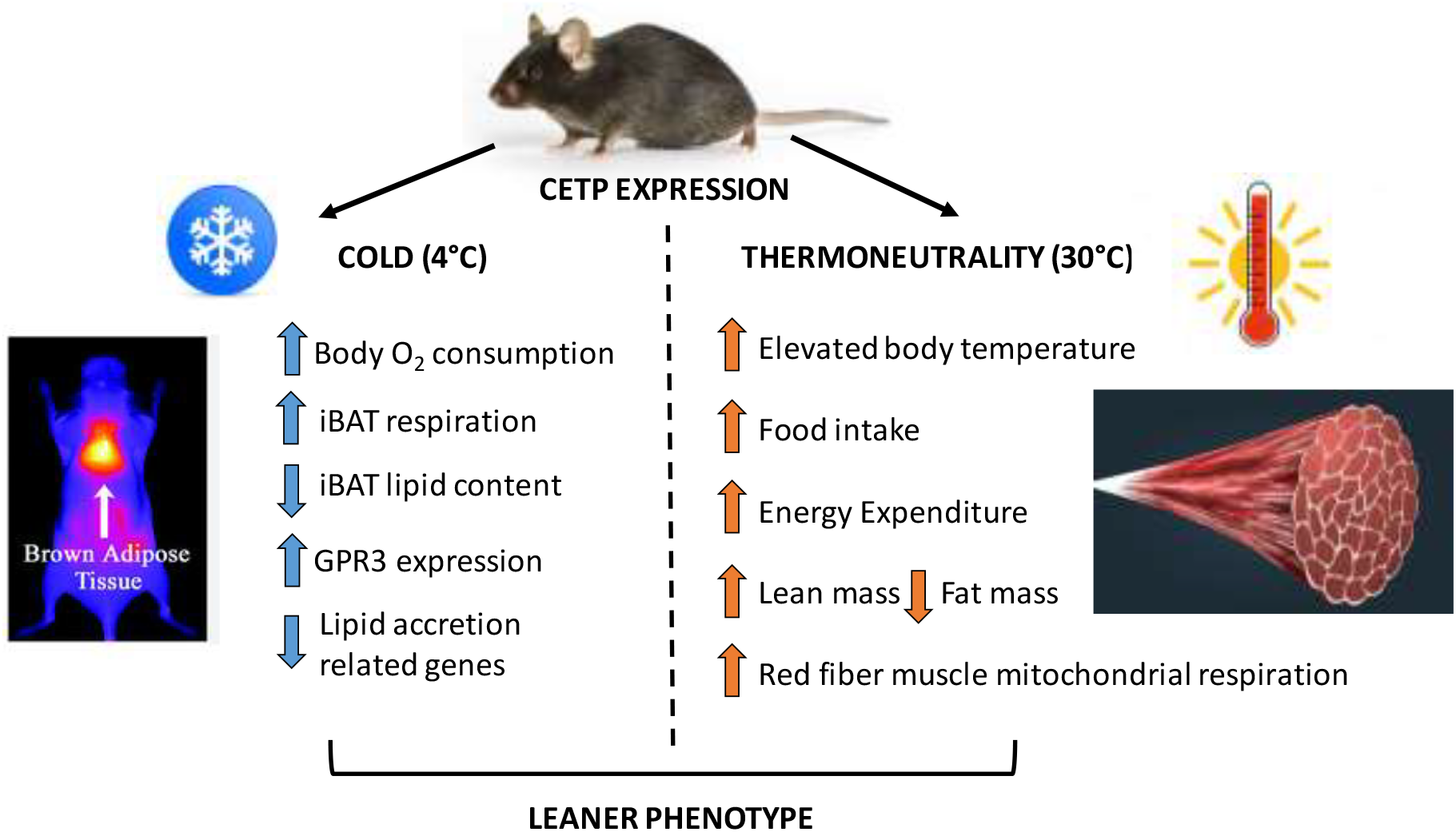

## INTRODUCTION

Cholesteryl ester transfer protein (CETP) plays an important role in lipid metabolism. Plasma CETP activity results in reduced HDL-cholesterol and, eventually, increased LDL-cholesterol concentrations, thus modulating the risk of cardiovascular diseases (Bruce and Tall 1995, Oliveira and Raposo 2020). Beyond its role on decreasing HDL-cholesterol, CETP seems to play other new functions also relevant for the cardiometabolic context. CETP may have tissue-specific HDL-independent local effects such as anti-inflammatory (Venancio et al 2016, Santana et al 2021, Rentz et al, 2023) and antioxidant (Dorighello et al 2022, Lazaro et al 2023). The body fat mass seems to be also a metabolic feature affected by the presence of CETP. When exposed to diet induced obesity and genetic hypertriglyceridemia, the expression of CETP reduces body fat mass and adipocyte area (Salerno et al 2007). Moreover, CETP expression in female mice on a chow diet decreases adiposity (Raposo et al 2021). Other metabolic protection mediated by CETP expression in female mice includes attenuation of obesity induced insulin resistance (Cappel et al 2013) and obesity induced decline in exercise capacity (Cappel et al 2015), as well as improvement of diet-induced fatty liver (Zhu et al 2022).

Environmental temperature is determinant of body metabolic rates and adaptive thermogenesis in mammals. Animal facilities are usually maintained at 22 ± 2°C, which is the threshold temperature for humans’ comfort and not necessarily the ideal for rodents (Keijer et al 2019). Thermoneutrality is particularly relevant for understanding the development of obesity, either for mice and men (Feldmann 2009). Rodents housing conditions at 18-22°C may cause chronical stress and food intake may increase 50-60% to keep body temperature (Cannon and Nedergaard 2009). Body heat can be generated by two main mechanisms: muscle contraction (physical activity or shivering) and non-shivering thermogenesis promoted by the brown adipose tissue (BAT) (Speakman 2013). Shivering, activated under acute cold exposure, may increase the oxygen consumption up to five times above the basal rate in men (Eyolfson et al 2001) and muscle fatigue occurs after longer periods of shivering (Wijers et al 2009). In this way, the non-shivering thermogenesis is especially important for cold acclimation. One essential mechanism of non-shivering thermogenesis during cold in mammals is the norepinephrine activation of BAT lipolysis that leads to mitochondrial uncoupling mediated by UCP1 and heat production (Cannon and Nedergaard 2017). On the other hand, in thermoneutrality (28-30°C for mice), UCP-1 dependent BAT activity is shut down, as demonstrated by similar oxygen consumption responses to norepinephrine (NE) in wild-type and UCP1 knockout mice (Cannon and Nedergaard 2011).

The leaner phenotype observed in CETP expressing female mice raised under standard conditions of 22ºC room temperature (moderate cold exposure) was associated with enhanced adipose lipolysis, body energy expenditure and brown adipose tissue activity (oxygen consumption and temperature) compared to CETP non-expressing littermates (Raposo at al 2021). Thus, in this study we aimed at investigating whether CETP expression modulate body temperature, composition, and metabolism when acclimated to cold (4°C) or under thermoneutrality (30°C).

## MATERIAL AND METHODS

### Experimental Animals

Two independent strains of CETP transgenic mice were used: **hCETP**, heterozygous CETP transgenic mice expressing a human natural promoter-driven CETP transgene (B6.CBA-Tg(CETP)5203Tall/J, RRID:IMSR_JAX:003904, Jiang et al., 1992) and **sCETP**, overexpressing simian CETP cDNA under the control of the metallothionein promoter (C57BL/6-Tg(CETP)UCTP20Pnu/J, RRID:IMSR_JAX:001929). Both mice line were purchased from The Jackson Laboratory (Bar Harbor, ME, USA) and crossbred with C57BL/6JUnib mice from the State University of Campinas Animal Center (RRID:MGI:7264953) for more than 10 generations. Mice were genotyped by PCR using tail tip DNA according to the Jackson Laboratory protocols. The non-transgenic (NTg) littermates were used as controls. Only female mice (aged 4-5 months) were used in this study to further investigated a phenomenon described only in females, a leaner phenotype (Raposo et al., 2021). Mice were maintained under controlled room temperatures (standard: 22 ± 2°C; cold exposure: 4 ± 1°C or thermoneutrality: 30±1°C), in a 12 h light/dark cycle, 15 cycles of air change per hour, with free access to filtered water and rodent standard chow diet (AIN93) (Nuvital CR1, Colombo, Brasil). The animal protocols were approved by the State University of Campinas Committee for Ethics in Animal Research (CEUA/UNICAMP, protocol #3870-1 and #5122-1/2019) and are in accordance with the Ethical Principles of the National Council for the Control of Animal Experimentation (CONCEA, Brazil) and the ARRIVE research guidelines for the use of Laboratory Animals (https://arriveguidelines.org). At the end of the experimental periods, mice were anesthetized with isofluorane inhalation and exsanguinated for blood and organs collection.

### Cold exposure

Female Mice (5 month-old) were exposed to cold (4 ºC) for 7 days, singly caged, in a 12 h light/dark cycle, with free access to water and chow diet. Body temperature (rectal temperature) was measured every 2 hours during the first 6 hours and once a day thereafter. No bedding and no food was offered during the first 6 hours. Food intake and body weight were evaluated 3 times during the 7-day experimental period.

### Thermoneutrality

Female Mice (4 month-old) were maintained at 30°C (2 heaters equally distributed in the room) for 7 days in a 12 h light/dark cycle, with free access to water and chow diet (2 mice/cage). Humidity was maintained at ∼60% with the constant use of a room humidifier. Food intake, body weight and rectal temperature were measured daily.

### Body Temperature

Body (rectal) temperature was measured using a thermometer with RET-3 rectal probe for mice (Physitemp, Clifton, NJ, USA) and regions of interscapular brown adipose tissue and tail temperature using a thermosensitive camera FLIR T450sc (FLIR Systems, Inc. Wilsonville, USA).

### In vivo Respirometry

In vivo respirometry was performed in an Oxylet System (Pamlab Harvard Apparatus, Barcelona, Spain). Mice were acclimated to the chamber for 24 hours and O_2_ and CO_2_ were measured during the next 24 hours. During the 48 hours in the chamber, mice had free access to food and water. The software Metabolism v2.2.01 was used to calculate total energy expenditure (EE). In separate experiments, oxygen consumption (VO2) was also evaluated in response to intraperitoneal injection of isoproterenol (10 mg/kg BW) in anesthetized mice (ketamine, 100 mg/kg and xylazine, 10 mg/kg).

### Carcass composition

The eviscerated carcasses were dehydrated at 65°C until body weight stabilized (dry weight). The total fat was then extracted with petroleum ether (LabSynth, SP, Brazil) in a Soxhlet extractor for 72 h. The fat mass is calculated by the weight difference of the dehydrated carcass before and after lipid extraction. Lean mass corresponds to the weight of dehydrated and delipidated carcasses.

### Brown adipose tissue histology

Samples were fixed with 4% paraformaldehyde during 24 h at 4°C, washed with phosphate buffered saline and preserved in ethanol 70%. Samples were then dehydrated in ethanol, and transferred to xylene solution for embedding in paraffin. Five μm longitudinal sections were cut in a Leica microtome. Slides were placed at 70°C to remove paraffin. Sections were stained with eosin-hematoxylin and digital images were captured under 40X objective lens in an optical microscope (Olympus BX51, camera Olympus U-TVO.63XC). The images were quantified using the ImageJ software (U.S. National Institutes of Health, Bethesda, MD, USA, http://imagej.nih.gov/ij). The percentage of fat present in the tissue was quantified using the method of Parlee and colleagues (2014). Lipid content was determined with the automated mode after binary transformation within fixed areas/section, five sections/mouse.

### Real-time polymerase chain reaction (RT-PCR)

Tissues were disrupted using stainless steel beads of 5 mm diameter in a Tissue Lyser LT (Qiagen, Venlo, The Netherlands). The resulting tissue homogenates were purified using the RNeasy Plus Mini Kit (Qiagen, Venlo, The Netherlands). The total RNA was quantified using a NanoDropTM 2000/2000c spectrophotometer (ThermoFisher Scientific, Waltham, MA, USA) and 1 μg was reverse transcribed using the High-Capacity cDNA Reverse Transcription Kit (ThermoFisher Scientific, Waltham, MA, USA) and a GeneAmp PCR System 9700 device (Applied Biosystems, Foster City, CA, USA). Reverse transcription was performed in four steps: 25ºC for 10 min, 37ºC for 120 min, 85ºC for 5 min. Target gene amplification was carried out using Fast SYBR Green Master Mix (ThermoFisher Scientific, Waltham, MA, USA) or QuantiNova SYBR Green PCR kit (Qiagen, Venlo, The Netherlands). cDNA (200 ηg) and forward and reverse oligonucleotide primers (300 ηM) (Table 1) were used for the reaction carried out in the 7500 Fast Real-Time PCR System (Applied Biosystems, Foster City, CA, USA), following the steps: 95ºC for 20 s, and 40 cycles of denaturation at 95ºC for 3 s and annealing/extension at 60ºC for 30 s. Gene expression was normalized using beta-actin as internal control, and the relative abundance of mRNA was quantified using the threshold cycle method (ΔΔCT).

**Table 1.**
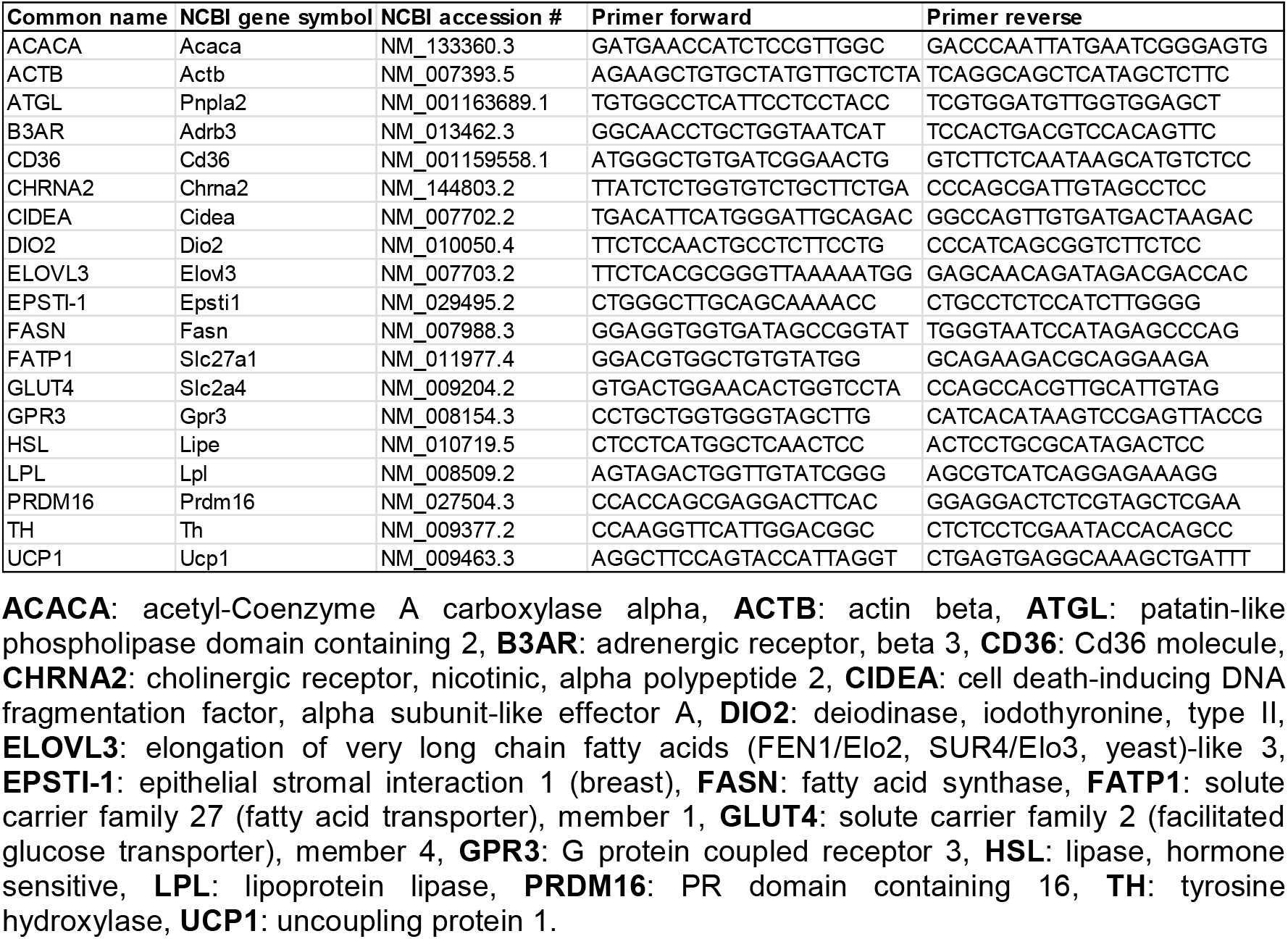
Primers for RT-PCR.

### High-Resolution Oxygraphy of tissue biopsies

Skeletal muscles (gastrocnemius and soleous) were freshly dissected and prepared in BIOPS medium supplemented with 10 mM Ca-EGTA buffer (2.77 mM CaK_2_EGTA + 7.23 mM K_2_EGTA, pH 7.1), 0.1 μM free calcium, 20 mM imidazole, 50 mM K^+^/MES, 0.5 mM dithiothreitol, 7 mM MgCl_2_, 5 mM ATP, and 15 mM phosphocreatine. Tendons and fascia were removed, and muscles were cut into small fragments or fiber bundles, each weighing between 2.5 and 5 mg. Muscles from each hindlimb were processed and analyzed separately as independent samples. Following previously established protocols (Busanello et al., 2017; Kuznetsov et al., 2008; Pesta and Gnaiger, 2015), the muscle fiber bundles were weighed, permeabilized in BIOPS medium containing saponin (50 μg/mL) with gentle stirring for 30 minutes at 4°C, and then washed three times in MiR05 (10 mM KH_2_PO_4_, 3 mM MgCl_2_, 500 μM EGTA, 60 mM lactobionic acid, 20 mM taurine, 110 mM sucrose, 1 g/L BSA, and 20 mM HEPES, pH 7.1) at 4°C. Liver slices were fragmented in a tissue chopper and processed as described above for muscles. Oxygen consumption rates (OCR) were measured using high-resolution oxygraphy (OROBOROS Oxygraph-2k, Innsbruck, Austria) in a 2 mL chamber. Samples (2.5–5 mg) were incubated at 37°C in MiR05 containing 10 mM pyruvate, 5 mM malate, and 10 mM glutamate. After measuring the baseline OCR, various respiratory states were induced by additions of 400 μM ADP, 0.2 × 3 (0.6) μM FCCP, 1 μM rotenone, 5 mM succinate, 1 μM antimycin A, oligomicin (1 µg/mL). Interscapular brown adipose tissue (iBAT) explants (3–4 mg), sliced with a scalpel, were incubated in DMEM containing permeable substrates (25 mM glucose, 2 mM glutamine, 1 mM pyruvate) supplemented with 4% fatty-acid-free BSA (w/v). iBAT oxygen consumption rates were monitored in the OROBOROS Oxygraph-2k in this basal condition. Recordings after 10 min of the experiment, when respiration rates were stable, were analyzed and normalized by tissue weight.

### Citrate synthase activity

The reaction solution consisted of 50 mM Tris, 0.1% Triton X-100, 50 µM acetyl-CoA, and 100 µM 5,5′-dithiobis(2-nitrobenzoic acid) (DTNB), with the pH adjusted to 8.0. Mouse soleus muscle extracts (0.5 µg/mL) were preincubated for 1 minute before initiating the reaction by adding 250 µM oxaloacetate. A blank sample without oxaloacetate was also included for comparison. The increase in absorbance at 412 nm was monitored over time using a microplate reader (PowerWave XS 2, BioTek Instruments, Winooski, VT, USA). This increase in absorbance indicates the formation of thionitrobenzoic acid, resulting from the citrate synthase-catalyzed production of citrate and CoA-SH. A molar extinction coefficient of 13.6/mM/cm was employed for calculations after determining the optical path length.

### Statistical Analyses

The data are expressed as mean ± standard error (SE) and were analyzed using GraphPad Prism®7 software. Outliers were tested using the Grubb’s test (alpha= 0.05) and normal distribution of the data was tested with Shapiro-Wilk test. Student’s t-test was employed to compare the two experimental groups and the level of significance was set at p≤ 0.05.

## RESULTS

Two independent CETP transgenic mice lines, expressing human (hCETP) or simian (sCETP) CETP transgenes were exposed to cold (4ºC). Rectal temperatures did not differ in CETP expressing mice compared to NTg either acutely (**Fig. 1 A,D**) or chronically, along 7 days of cold exposure (**Fig. 1 B,E**). During cold exposure, CETP expression did not alter food intake (**Fig. 1 C,F**), body weight (**Fig. 2 A,C**) and mass of visible adipose depots (**Fig. 2 B,D**). In sCETP group, we observed that the interscapular BAT mass (iBAT) was significantly increased (15%) when compared to NTg (**fig. 2 B,D**).

**Figure 1.**
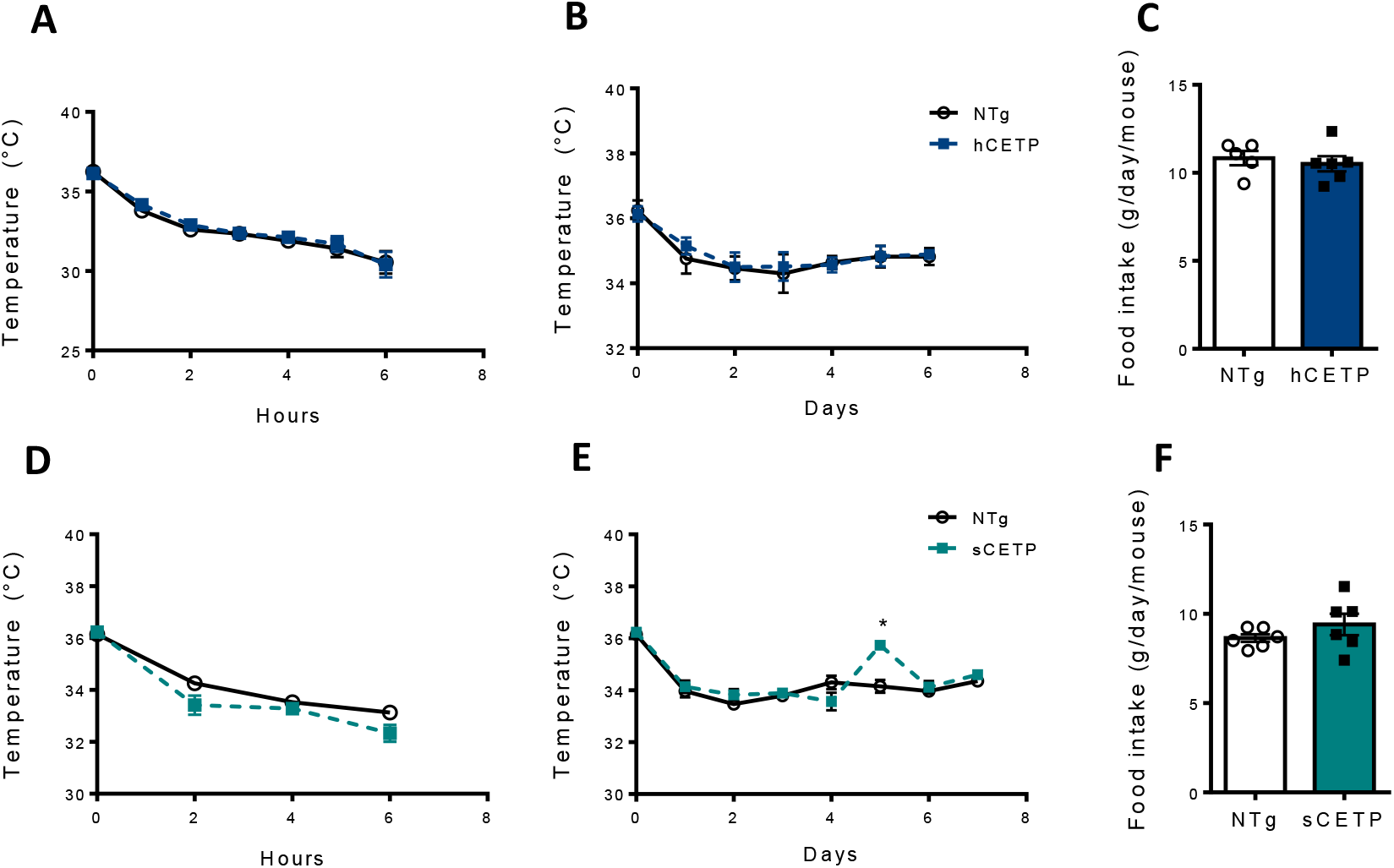
Body (rectal) temperature during acute (**A,D**) and chronic (**B,E**) cold exposure (4ºC) and food intake (**C,F**) of human CETP (hCETP) (**A,B,C**) and simian CETP (sCETP) (**D,E,F**) female mice compared to their non-transgenic (NTg) controls. Mean ± SE (n=5-6). Student’s t test, * p ≤ 0.05.

**Figure 2.**
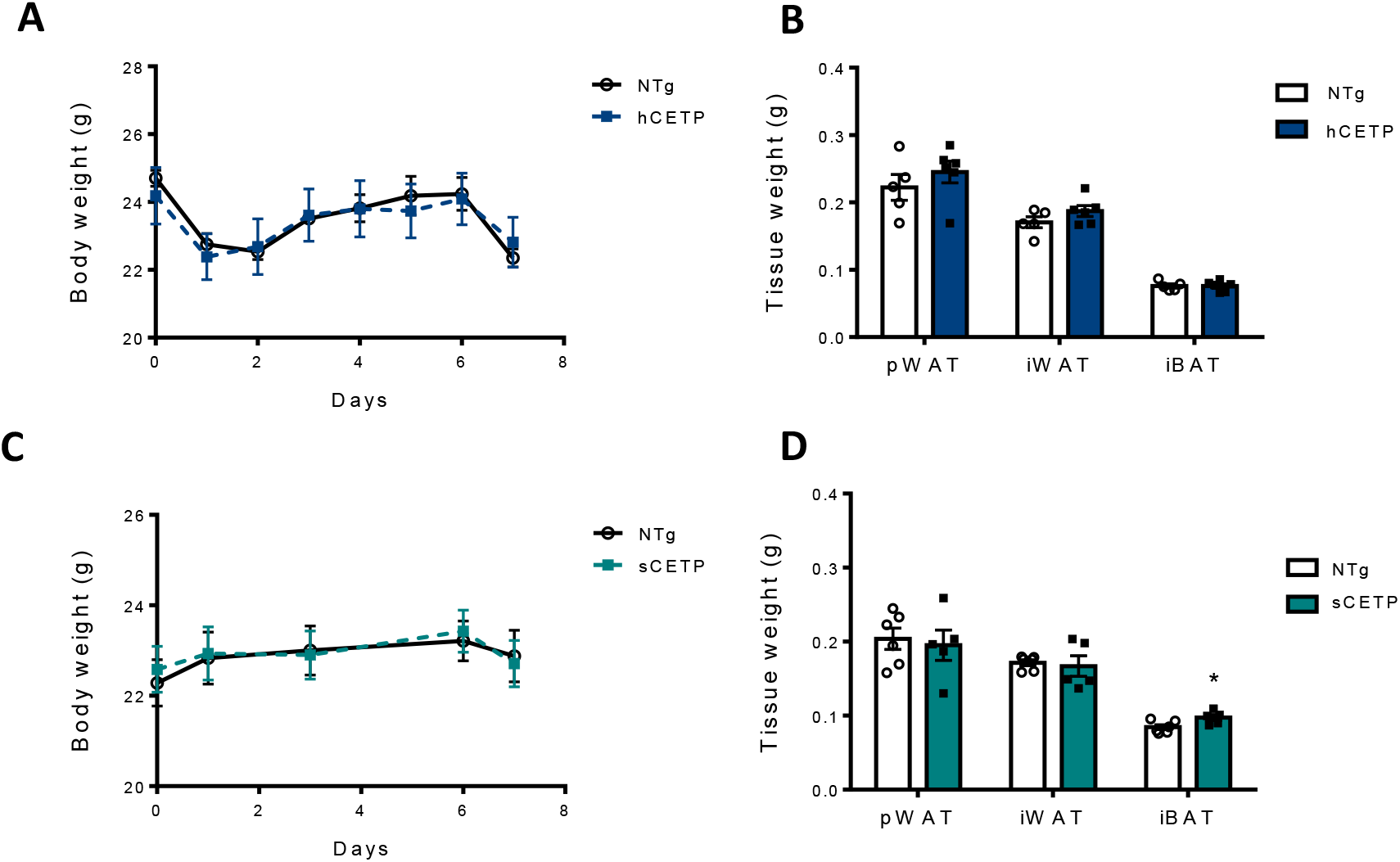
Body weight (**A,C**) and mass of perigonadal (pWAT), inguinal (iWAT) white adipose and interscapular brown adipose (iBAT) tissues (**B,D**) during chronic cold exposure (4ºC) of hCETP (**A,B**) and sCETP (**C,D**) female mice compared to their non-transgenic (NTg) controls. Mean ± SE (n=5-6). Student’s t test, * p ≤ 0.05.

We then hypothesized that CETP mice, although leaner before the cold exposure and thus with less insulation, are able to maintain their body temperature during cold exposure similarly to NTg due to a higher capacity to increase BAT activity. To test that, we submitted mice to respirometry in response to a norepinephrine analogue, isoproterenol, administration. These analyses were performed before (housing at 22°C) and after 7 days of cold exposure (**Fig. 3 A,B**). At both housing temperatures (22 and 4°C), the body oxygen consumption rates in response to isoproterenol were higher in CETP group compared to NTg. While an increment of 21% was seen at 22ºC (**Fig. 3 A**), after 7 days of cold exposure, the CETP group had a 41% increased oxygen consumption rate compared to NTg (**Fig 3 B**). This indicates that cold exposure further increased iBAT metabolism in CETP expressing compared to non-expressing mice. In agreement with this functional in vivo test, cold exposed CETP mice showed increased basal oxygen consumption rate measured ex vivo in isolated iBAT biopsies (**Fig 3 C**) and reduced iBAT lipid content (**Fig. 3 D,E**).

**Figure 3.**
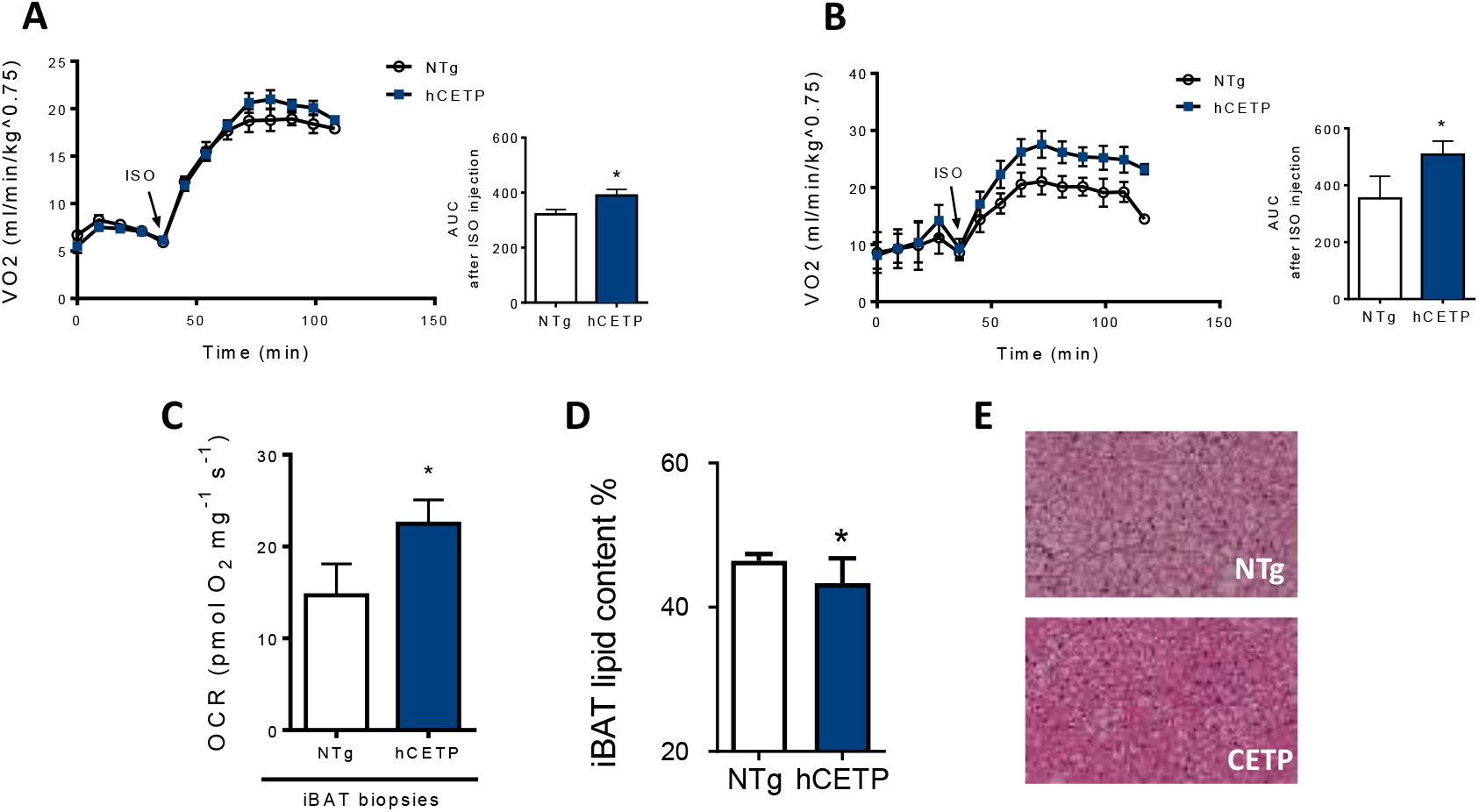
In vivo body oxygen consumption after isoproterenol administration before (22ºC) (**A**) and after 7 days at 4ºC (**B**) in hCETP female mice and non-transgenic (NTg). AUC: area under the curves after isoproterenol i.p. injection (ISO: 10 mg/kg BW). In situ iBAT biopsies basal oxygen consumption rates (OCR) (**C**) and lipid content (**D**). Representative histological iBAT images in 10x magnification (**E**). Mean ± SE (n=5-6 for VO2 and n=7-8 for OCR and histology). Student’s t test, *p ≤ 0.05.

In order to confirm depletion of lipid substrates in cold exposed CETP mice, and gain some mechanistic insight, we analyzed the expression of a panel of lipid metabolism controlling genes in two depots, iBAT and inguinal (subcutaneous) adipose tissue (iWAT) (**Fig. 4**). We found that CETP mice have a 50% increase above NTg mice levels of GPR3 (G protein-coupled receptor 3) expression in both adipose tissues. GPR3 is a constitutive receptor activated by intracellular lipolysis, cold induced and thermogenesis stimulant (Sveidahl Johansen et al, 2021). In addition, CETP adipose depots exhibited downregulation of several genes encoding proteins related to lipid accumulation (LPL, ELOV3, ACACA, FAS, CD36, FATP1, CIDEA and GLUT4).

**Figure 4.**
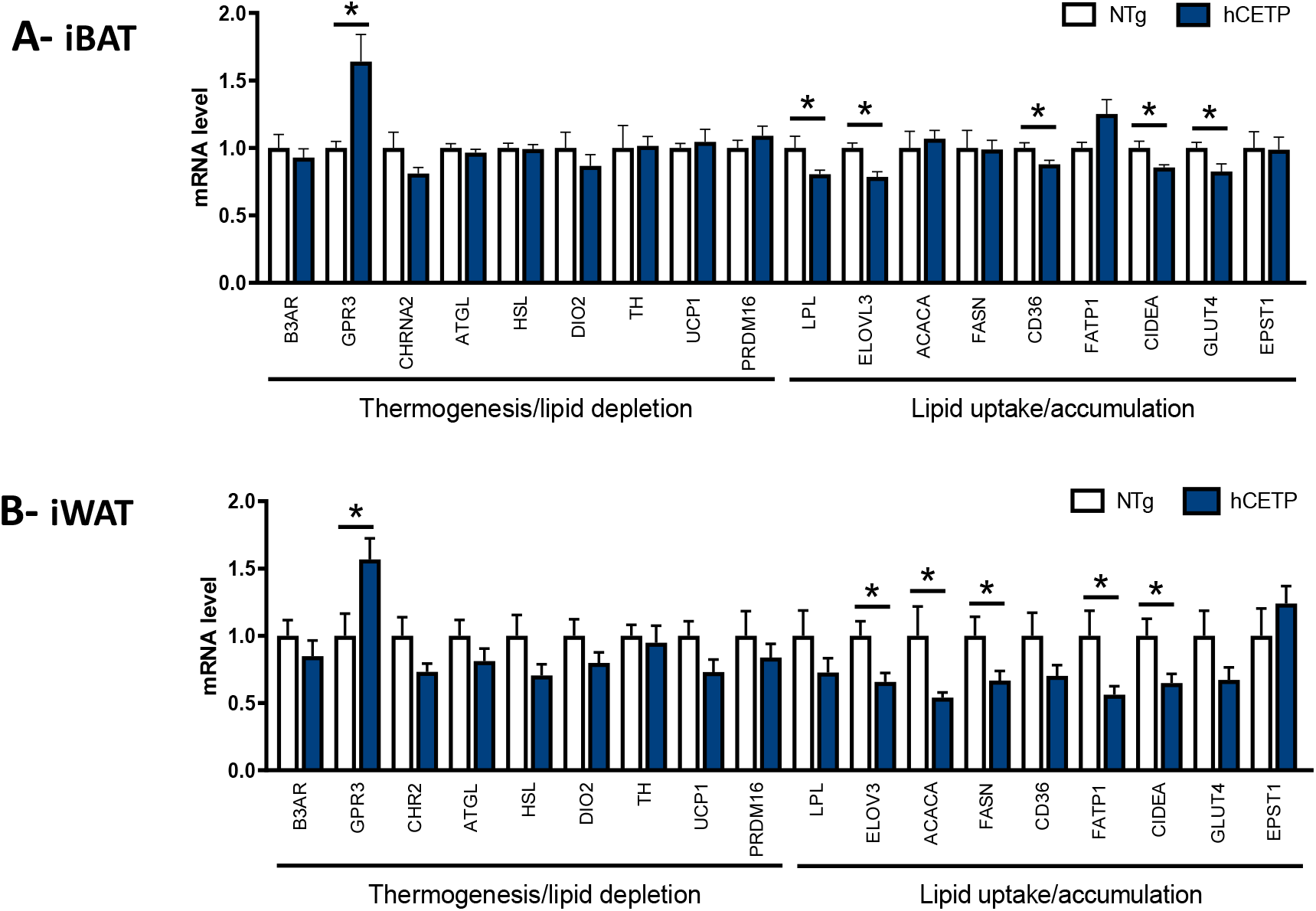
Relative mRNA levels of thermogenic and lipid metabolism related genes in iBAT (**A**) and iWAT (**B**) of hCETP female mice and non-transgenic (NTg) and after 7 days at 4ºC. Data normalized by beta-actin as internal control. Mean ± SE (n= 7-8). Student’s t test, *p ≤ 0.05. **ACACA** (*Acaca*): acetyl-Coenzyme A carboxylase alpha, **ATGL** (*Pnpla2*): patatin-like phospholipase domain containing 2, **B3AR** (Adrb3): beta-3 adrenergic receptor, **CD36** (*Cd36*): Cd36 fatty acid transporter, **CHRNA2** (*Chrna2*) cholinergic receptor, nicotinic, alpha polypeptide 2, **CIDEA** (*Cidea*): cell death-inducing DNA fragmentation factor, alpha subunit-like effector A, **DIO2** (*Dio2*): deiodinase-2, **ELOVL3** (*Elovl3*): elongation of very long chain fatty acids, **EPSTI-1** (*Epsti1*): epithelial stromal interaction 1, **FASN** (*Fasn*) fatty acid synthase, **FATP1** (*Slc27a1*): solute carrier family 27, fatty acid transporter 1, **GLUT4** (*Slc2a4*): solute carrier family 2 member 4, **GPR3** (*Gpr3G*): G protein coupled receptor 3, **HSL** (*Lipe*): hormone sensitive lipase, **LPL** (*Lpl*): lipoprotein lipase, **PRDM16** (*Prdm16*): PR domain containing 16, **TH** (*Th*): tyrosine hydroxylase, **UCP1** (*Ucp1*): uncoupling protein 1.

Next, we checked whether CETP expression would affect body composition and metabolism under thermoneutrality, a condition where BAT activity is inhibited. Thus, CETP mice and their NTg littermates were maintained at 30ºC for 7 days. Under this condition, CETP mice presented higher body temperatures, both core rectal and surface tail temperatures, but iBAT surface temperature was similar between groups, as expected (**Fig. 5**). Body weight did not vary between groups (**Fig. 6 A**), while food intake was increased in CETP mice during thermoneutrality (**Fig. 6 B**). The mass of white adipose visible depots were similar between groups and, as expected, BAT mass and lipid content did not differ between groups (**Fig. 6 C,D)**. However, we observed significant changes in carcass composition, with increased lean mass and decreased fat mass in CETP compared to NTg mice maintained at 30ºC (**Fig. 6 E**).

**Figure 5.**
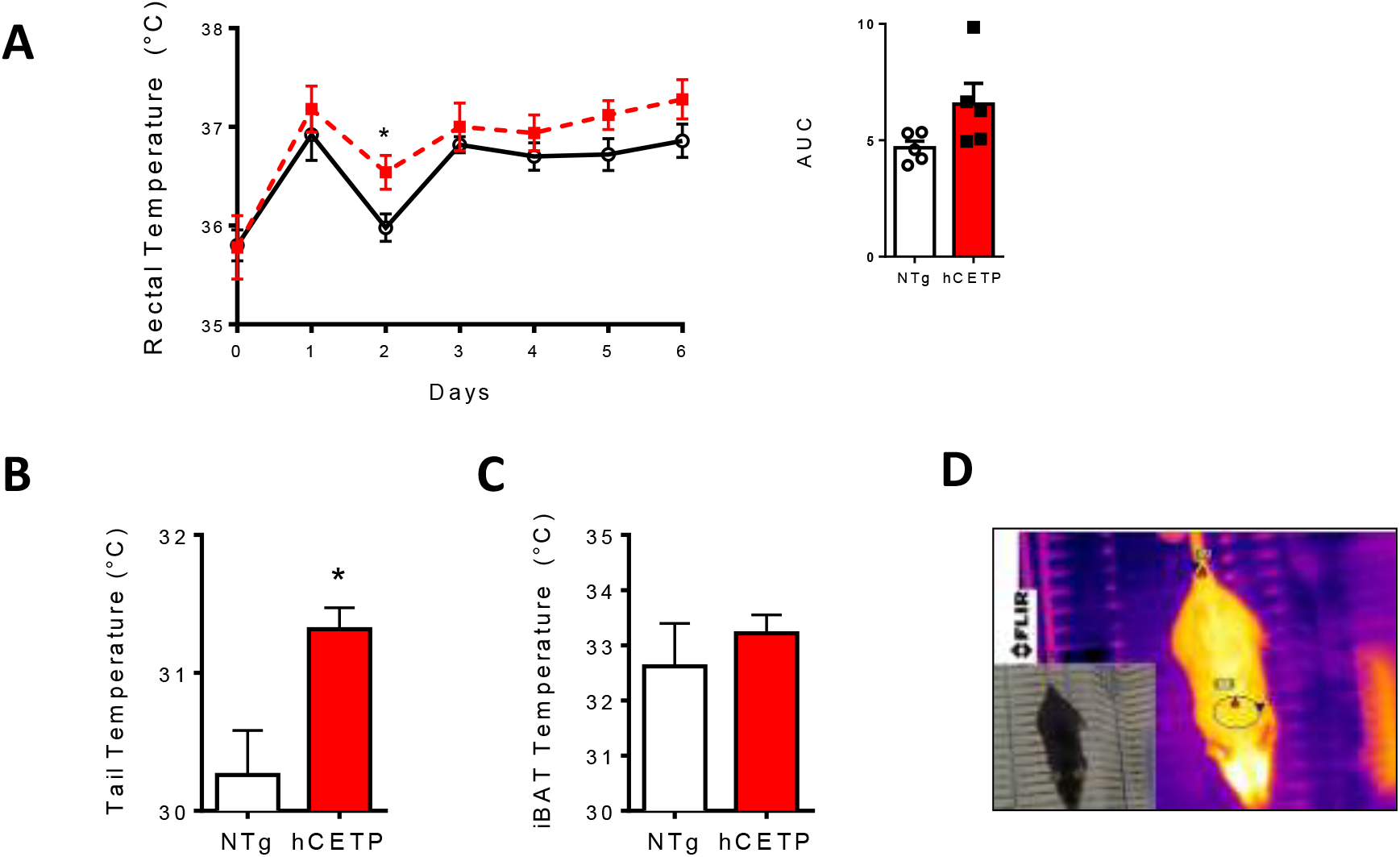
Female hCETP mice present higher core (rectal) (**A**) and surface (tail) (**B**), but similar iBAT (**C**) temperatures compared with NTg mice under thermoneutrality (30ºC) for 7 days. Representative image of thermographic picture to estimate iBAT and tail temperatures (**D**). Image red and blue triangles indicate maximum and minimum temperatures, respectively, in each region. Mean ± SE (n=5-6). Student’s t test, * p ≤ 0.05.

**Figure 6.**
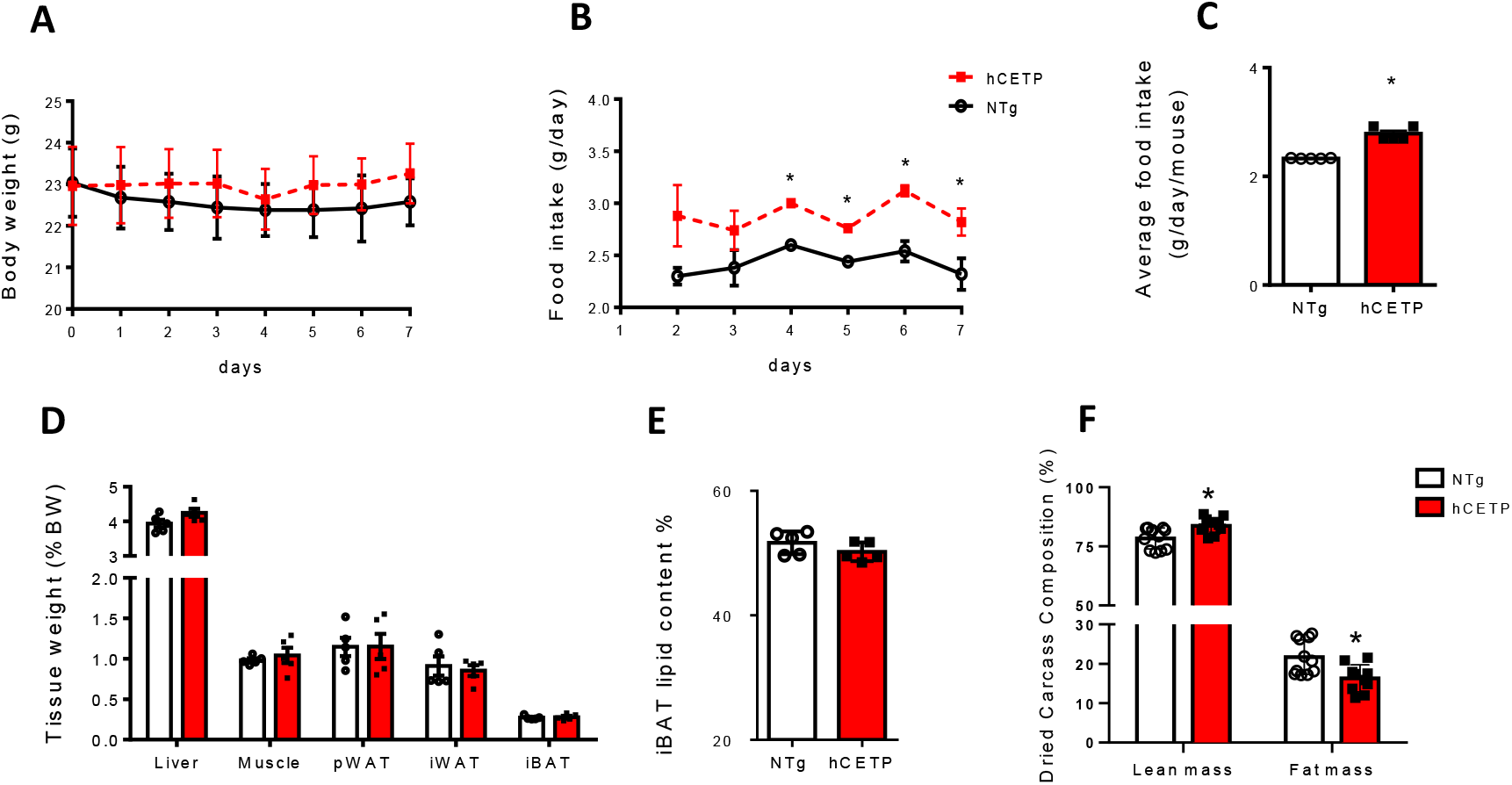
Female hCETP mice present increased food intake and lean mass and decreased fat mass at thermoneutrality. Body weight (**A**), food intake (**B, C**), tissues mass (**D**), iBAT lipid content (**E**) and dried carcass composition (**F**) after 7 days at 30ºC. Mean ± SE (n=5-6). Student’s t test, * p ≤ 0.05.

Higher body temperature and food intake together with reduced fat mass can only be explained by elevated body metabolic rate. In fact, respirometry analyses showed that CETP mice presented higher energy expenditure (EE) during the light period of the day (**Fig. 7**). In order to find out which tissue would contribute more importantly to the higher EE in CETP mice, we performed ex vivo high resolution oximetry in tissue biopsies. In the thermoneutrality condition, we observed that soleus muscle (**Fig. 8 A**), but not gastrocnemius or liver (**Fig. 8 B,C**) of CETP mice showed increased mitochondrial respiration rates, under basal and phosphorylating (ADP addition) conditions, as well as at maximum FCCP-induced uncoupled respiration. The amount of mitochondria in the soleus muscle seems not to be different between groups as indicated by the activity of the mitochondrial enzyme citrate synthase (0.378±0.024 vs 0.331±0.042 mU mg^-1^, for NTg and CETP, respectively, p=0.35) and by the ratio FCCP/basal respiration rates which correlates with cytochrome oxidase activity (1.75±0.096 vs 1.92±0.149 for NTg and CETP, respectively, p=0.35). This suggests that mitochondrial content is similar in both groups and increased respiration rates in CETP mice may be attributed to the tissue substrates availability or the presence of other stimulating factors.

**Figure 7.**
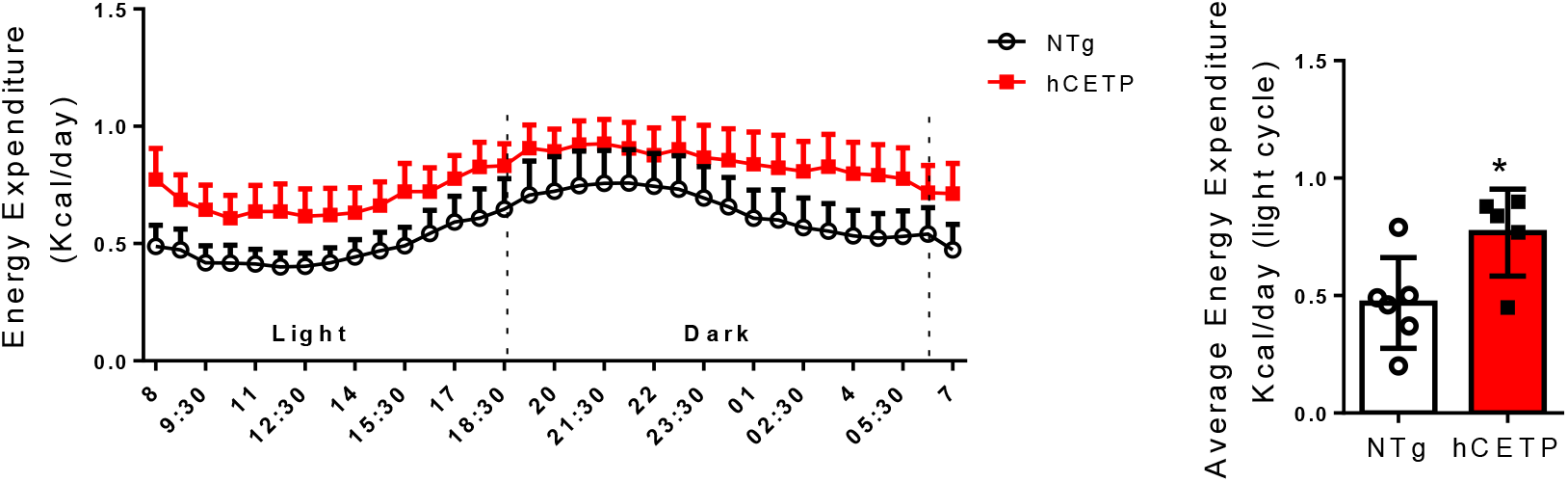
Female hCETP mice exhibit higher body metabolic rate measured as energy expenditure (EE) during the resting light cycle of the day under thermoneutrality (**A, B**). Mean ± SE (n=5-6). Student’s t test, * p ≤ 0.05.

**Figure 8.**
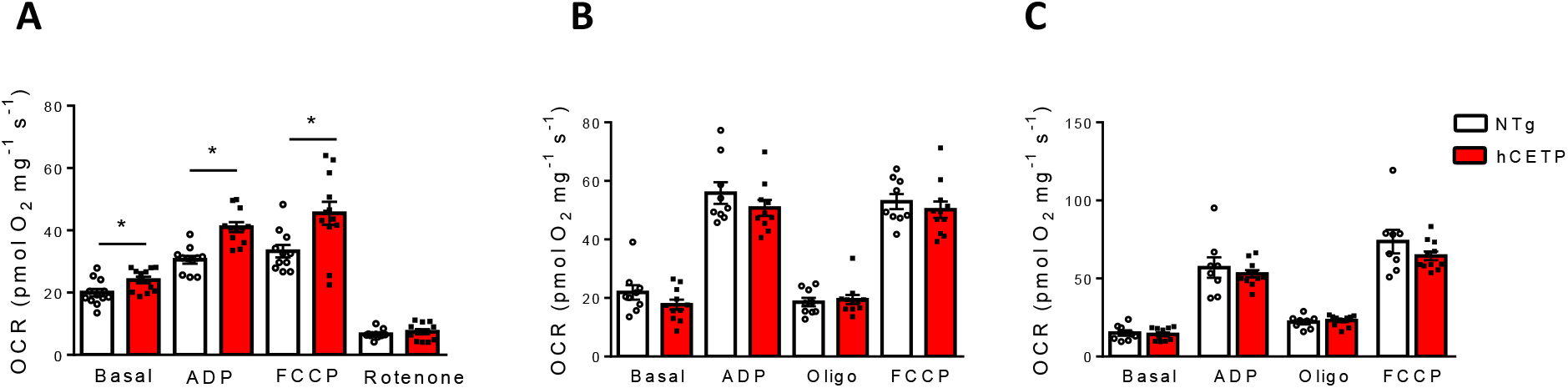
Increased respiration (oxygen consumption rates, OCR) in the soleus muscle (**A**), but not in gastrocnemius (**B**) or liver (**C**) of female hCETP mice acclimated to thermoneutrality. Respiration was evaluated in the MiR05 medium at 37ºC containing 10 mM glutamate plus 5 mM malate as substrates. Respiration conditions are: basal (no additions), phosphorylating (ADP, 400 μM), maximum respiration (FCCP, 0.6 μM), resting respiration (oligomycin, 1 μg/mL) and non-mitochondrial respiration (rotenone, 1 μM). Mean ± SE (n=10-12). Student’s t test, * p ≤ 0.05

## DISCUSSION

Previous studies have shown that susceptibility to obesity differs depending on the genetic background and housing temperatures of experimental models. Cold exposure/acclimation increases BAT volume and activity in human adults (Saito et al 2009, Enerbäck et al 2010, Blondin et al 2014) and increases body oxygen consumption rate and reduces obesity in rodents (Kajimura et al 2008, Lowell and Spiegelman 2000). We have previously reported that under chow diet and housing temperature at 22ºC, CETP expression increases white and brown adipose lipolysis, BAT oxygen consumption and body metabolic rate, resulting in reduced adipose tissue mass as compared to NTg mice (Raposo et al 2021). Here, we show that these 22ºC acclimated CETP mice when exposed acutely or chronically to cold (4ºC) are able to maintain their body temperature similarly to NTg, even though they had less insulation (fat mass) prior to cold. The explanation for that is linked to their higher capacity to further increase metabolic rate induced by cold, shown by: 1) higher isoproterenol induced body oxygen consumption, 2) higher BAT consumption of lipid substrates and 3) higher BAT oxygen consumption rates in CETP mice. In small mammals, BAT is the main contributor to adaptive non-shivering thermogenesis that occurs under cold acclimation (Cannon and Nedergaard 2011). Although the contribution of other tissues, such as liver and white adipose tissue to adaptive thermogenesis in humans is uncertain (Lowell and Spiegelman 2000), other findings identified liver as a relevant site for cold adaptation (Simcox et al 2017), since liver provides acylcarnitines as a fuel for BAT non-shivering thermogenesis in cold-exposed mice.

Prolonged exposure to cold induces the development of brown-like adipocytes embedded in white adipose tissue (WAT), a phenomenon called “browning” (Petrovic et al 2010, Vegiopoulos et al 2010, Okamatsu-Ogura et al 2018). To understand better the increase in BAT activity and to check a putative browning of white adipose tissue in CETP mice, we screened for key lipid metabolism and thermogenesis related gene expression pattern. The most noteworthy feature observed in both adipose depots of CETP mice is the marked increase in the expression of GPR3, a newly described receptor activated by intracellular lipolysis products, essential for cold induced thermogenesis (Balaz and Wolfrum 2021, Sveidahl Johansen et al 2021). Lack of GPR3 in mice leads to impaired BAT thermogenic function and late onset obesity (Godlewski et al 2015), while activation of GPR3 inhibits obesity and liver pathogenesis (Dong et al 2024). Thus, the observed CETP cold induced higher metabolism is likely associated with this higher expression of GPR3. In the previous study (Raposo et al 2021), we demonstrated increased lipolysis in WAT and BAT of CETP mice, what explains the enhanced GPR3 gene expression (and likely activity) which is driven by lipolysis (Sveidahl Johansen et al 2021).

Although we found no alterations in UCP1 gene expression in iWAT and iBAT of cold exposed CETP mice, it does not mean that UCP1 is not more active in these tissues. The availability of lipolysis-derived non-esterified fatty acids is sufficient to increase UCP1 activity resulting in heat production and energy dissipation (Alberici and Oliveira, 2022). Other mitochondrial uncoupling mechanisms elicited by increased fatty acids availability, such as activation of mitochondrial ATP sensitive potassium channel (Alberici et al 2011), may not be excluded. Interestingly, several lipid accumulation related genes were downregulated in iBAT and iWAT of CETP mice, which may represent an adaptive response to cold. This also explains in part the leaner phenotype of female CETP mice.

The possible reasons why CETP mice have increased lipolysis compared to their NTg counterparts were researched previously in mice maintained at 22ºC (Raposo et al 2021). Increased cholesterol content in adipocyte membranes, such as observed in iBAT of CETP mice, may stabilize surface receptors such as ß3-adrenergic receptors that are crucial to signal lipolysis (Holm 2003). Elevation of bile acids are known activators of thermogenesis (Watanabe et al 2006). We did not find higher total circulating steady state levels of bile acids in CETP expressing mice, but increased bile acid turnover was shown by Cappel et al. (2013) in obese female CETP mice, stimulating BAT activity via TGR5 receptors and iodothyronine deiodinase 2 activity.

When acclimated to thermoneutrality, CETP expressing mice exhibit significant increases in body temperature, suggesting that the higher metabolic rate of CETP mice is a constitutive phenomenon in this model. Since core (rectal) and surface (tail) temperature are elevated, it seems that both heat generation and dissipation are increased in these mice. The higher body metabolic rates were confirmed by respirometry analyses showing higher energy expenditure during the light period of the day in CETP mice acclimated to thermoneutrality. Since BAT activity is inhibited in thermoneutral conditions, we searched for other metabolic active target tissues. Considering that CETP mice showed increased lean mass and decreased fat mass under thermoneutrality, we hypothesized that skeletal muscle could have a great contribution to keep higher body metabolism in CETP mice, not only because of a larger mass but mainly by elevated metabolism. We then determined oxygen consumption rates in mixed fibers muscle gastrocnemius, red fiber muscle soleus and also in liver, a tissue that exhibits high metabolic rates. We observed that soleus muscle, but not gastrocnemius or liver of CETP mice showed increased mitochondrial respiration rates, under basal and phosphorylating conditions, as well as at maximum induced mitochondrial respiration rates. The increased mitochondrial respiration rates is not related to the tissue mitochondrial content, but could be attained by a higher efficiency of the respiratory chain and/or increased endogenous substrate uptake and metabolism. In agreement with our findings, female mice expressing simian CETP fed with a high fat diet had improved exercise capacity due to increased muscle mitochondrial oxidation of malate/glutamate substrates (Cappel et al 2015). Although we did not investigate the underlying mechanisms of red fiber skeletal muscle elevated metabolism in CETP mice under thermoneutral environment, we could speculate that heat generated in resting skeletal muscles, uses the same muscle cellular machinery to regulate cytoplasmic calcium concentrations and contraction. Micro movements in [Ca2+] and associated changes in ATP turnover are key events for the process known as muscle-based non-shivering thermogenesis, coordinated by the plasma membrane, sarcoplasmic reticulum and the mitochondria (Launikonis and Murphy 2025).

It is also possible that other tissues contribute to the elevated body metabolism of CETP mice either under cold or thermoneutrality. We demonstrated previously that CETP-expressing macrophages (from mice acclimated to 22ºC) exhibited increased mitochondrial maximal respiration rates and spare respiratory capacity (Dorighello et al 2022). Thus, macrophage rich tissues from CETP mice could also possess higher mitochondrial metabolism.

In conclusion, leaner female CETP mice are able to maintain their body temperature during acute and chronic cold exposure due to their higher capacity to increase body metabolic rates and BAT activity above the control NTg mice. Under thermoneutrality (when BAT activity is reduced), female CETP mice exhibited elevated body temperature, food intake, energy expenditure, lean mass and red fiber muscle mitochondrial respiration rates. Thus, we propose that CETP expression confers a greater capacity of elevating body metabolic rates under distinct housing temperatures, what explain their leaner phenotype. These findings suggest that CETP expression levels in females should be considered as a new influence in the contexts of obesity and metabolic disorders.

## Data Availability Statement

Data supporting the findings of this study are available from authors upon reasonable request.

## Conflicts of Interest

The authors declare no conflict of interest.

## Authors’ contributions

JZC, HFR, CDCN, CML, MRS, APDC, PASN, performed experiments, analyzed data, interpreted results and prepared figures; HFR drafted the first version of the manuscript; LAV and AEV provided methodology and resources, interpreted results and revised the manuscript. HCFO conceived and designed research, methodology and resources, analyzed data and wrote and revised the manuscript. All authors read and approved the final manuscript.

## Funding

This work was supported by grants from Fundação de Amparo à Pesquisa do Estado de São Paulo (FAPESP #2013/07607-8 to HCFO, AEV and LAV; and #2017/17728-8 to HCFO and AEV) and Conselho Nacional de Desenvolvimento Científico e Tecnológico (CNPq #300937/2018-0 to HFCO) and FAPESP scholarships: #2019/02304-0 to JZC, #2015/17555-0 to HFR, #2019/13862-7 to CML and #2023/02270-7 to CDCN.

## Acknowledgments

We are especially grateful to Dr. Cesar Sartori and Dr. André Vieira from the Department of Structural and Functional Biology at the State University of Campinas for their support with the animal facility for thermoneutrality experiments.

## Notes

### Competing Interest Statement

The authors have declared no competing interest.

